# Characterisation of Geometric Variance in the Epithelial Nerve Net of the Ctenophore *Pleurobrachia pileus*

**DOI:** 10.1101/2020.06.30.178582

**Authors:** Amy Courtney, Jérémy Liegey, Niamh Burke, Madeleine Lowey, Mark Pickering

**Affiliations:** School of Medicine, University College Dublin, Ireland; School of Electrical & Electronic Engineering, University College Dublin, Ireland; UCD Centre for Biomedical Engineering, University College Dublin, Ireland

**Keywords:** ctenophore, nerve net, model organism, neuroscience

## Abstract

Neuroscience currently lacks a diverse repertoire of model organisms, resulting in an incomplete understanding of what principles of neural function generalise and what are species-specific. Ctenophores display many neurobiological and experimental features which make them a promising candidate to fill this gap. They possess a nerve net distributed across their outer body surface, just beneath the epithelial layer. There is a long-held assumption that nerve nets are ‘simple’ and random while lacking distinct organisational principles. We want to challenge this assumption and determine how stereotyped the structure of this network really is. We validated an approach to estimate body surface area in *Pleurobrachia pileus* using custom Optical Projection Tomography and Light Sheet Morphometry imaging systems. We used an antibody against tyrosylated *α*-tubulin to visualise the nerve net in situ. We used an automated segmentation approach to extract the morphological features of the nerve net. We characterised organisational rules of the epithelial nerve net in *P. pileus* in animals of different sizes and at different regions of the body. We found that specific morphological features within the nerve net are largely unchanged during growth. These properties must be essential to the functionality of the nervous system and therefore are maintained during a change in body size. We have also established the principles of organisation of the network and showed that some of the geometric properties are variable across different parts of the body. This suggests that there may be different functions occurring in regions with different structural characteristics. This is the most comprehensive structural description of a nerve net to date. This study also demonstrates the amenability of the ctenophore *P. pileus* for whole organism network analysis and shows their promise as a model organism for neuroscience, which may provide insights into the foundational principles of nervous systems.

## Introduction

### Diversity of Model Organisms in Neuroscience

Due to the technical challenges arising from the complexity of the nervous system, particularly in humans, a long standing approach in neuroscience research is to instead focus on simpler, experimentally tractable model organisms to explore fundamental principles of organisation. With many scientists working with the same model organism, many tools and resources are established, and this is an effective way to enrich our scientific understanding of specific biological processes. However, this comes at a cost in that the larger structural and functional diversity of animal nervous systems is ignored. This results in a lack of understanding of what principles of neural function generalise and what are species-specific and this in turn hinders our ability to extrapolate this knowledge to the human brain. Some researchers are now advocating for the field of neuroscience to expand its repertoire of model organisms to increase diversity and promote an understanding of nervous system evolution and function more broadly (Keifer & Summers, 2016; Striedter et al., 2014; Yartsev, 2017).

### The Importance of a Comparative Approach: Cnidaria and Ctenophora

The Cnidaria have been promoted as important models to fill this gap and many tools have been developed to promote their use in this arena (Bosch et al., 2017). A recent breakthrough saw the cnidarian *Hydra vulgaris* become the first animal in which the activity of all the neurons in a nervous system were simultaneously recorded while the animal was freely behaving (Dupre & Yuste, 2017). However, the Ctenophora have not received the same consideration despite possessing many of the same neurobiological and experimental advantages as the Cnidaria. From a nervous system evolution perspective, both lineages diverged prior to the origin of bilaterally symmetrical animals and thus both could represent the earliest extant forms of the first nervous systems. However, it has been suggested that the Ctenophora have evolved their nervous system independently to all other animals (Moroz et al., 2014). If this claim is substantiated, then these animals represent the only living example of convergent evolution of a nervous system. This has implications for their use as a model organism because studying traits that evolved independently in two groups provides insights into the fundamental properties of that system (Serb & Eernisse, 2008). On the other hand, if this claim turns out not to be true then these animals still represent the present-day form of the most ancestral nervous system and studying their nervous system will provide insights into the basic rules of organisation and function. From a technical perspective, both the Cnidaria and the Ctenophora possess a decentralised nerve net, this architecture is relatively simple, which makes it tractable for neuroanatomical studies. However, ctenophore neurons have been notoriously challenging to study and our understanding of the global morphology of their nerve net is incomplete.

### Epithelial Nerve net Organization in *Pleurobrachia pileus*

While detailed characterization of elements of the nervous system in ctenophores have been reported (Jager et al., 2011; Norekian & Moroz, 2019), a systematic description of the nervous system architecture has not yet been completed. The epithelial nerve net in the ctenophore *Pleurobrachia pileus* has been described as a polygonal lattice (Hernandez-Nicaise, 1973). The polygons are the spaces between the branched network of neurons when it is visualised in a wholemount preparation. The polygons are seemingly irregular in shape, size, density and orientation at different parts of the body, but no quantification or systematic network wide analysis has taken place. *Pleurobrachia bachei* has been described as possessing 5000-7000 neurons (Norekian & Moroz, 2019), but how these cells fit into the nerve net structure is not known. Each branch of the network which defines the sides of the polygon is comprised of 2-5 parallel neurites, with neuron cell bodies being found both along these branches and at the connecting nodes (Jager et al., 2011).

The term “nerve net” itself carries an implication of inherent “simplicity”, and a lack of distinct organizational principles. However, it is clear that distinct, predictable organization is evident in specific parts of the nerve net in *Pleurobrachia*. The juxtatentacular nerve (JTN) is a structure found along the tentacular plane of the animal extending from the aboral organ, into the tentacular sheath and is continuous with the base of the tentacle. It also extends from the opening of the tentacular sheath to the mouth (Jager et al., 2011). This structure is a condensation of neurites which is continuous with (and can be considered an element of) the epithelial nerve net. The function of the JTN is unclear. It was proposed that it is involved in propagating sensory information from the aboral organ, mouth, body surface and tentacles to control feeding, swimming and escape responses (Norekian & Moroz, 2019), although this is contrary to experiments performed by Moss & Tamm (1993) in which lesioning of this structure did not prevent a feeding response.

The presence of organised structures within the nerve net raises two specific but related questions. First, if the structure of the JTN is predictable and repeatable between individual animals, does the nerve net as a whole also look the same between animals? This would imply the existence of organizational rules which make the geometry and structure of the nerve net as a complete system predictable to a degree. The second question follows from this. If one part of the nervous system (the JTN) displays regional structural specialization, it implies that local differences can exist in the organizational rules governing the structure of the nerve net. If that’s the case for the JTN, could other local differences produce regional specialization in other regions of the nerve net?The JTN is the most obvious and clear case of regional specialization, but does it exist throughout the nerve net?To address these questions, we considered the problem that animal growth poses for the organization of a nervous system. This problem is particularly pronounced for animals with a nerve net closely aligned to the body surface. As the animal grows, and surface area increases, the structure of the nerve net must necessarily change, while the overall function of the nervous system (i.e. sensing and responding to the internal and external environment) must remain relatively unchanged. By determining which features of nerve net geometry remain constant, and which features change, we can gain insight into which morphological features might be important for that function. For example, as the body grows, the nerve net polygons might enlarge proportionately, suggesting the number of polygons is an important characteristic. Alternatively, the number of polygons may increase to occupy the increased area, suggesting the importance of maintaining polygon size (Figure 1).

**Figure 1.**
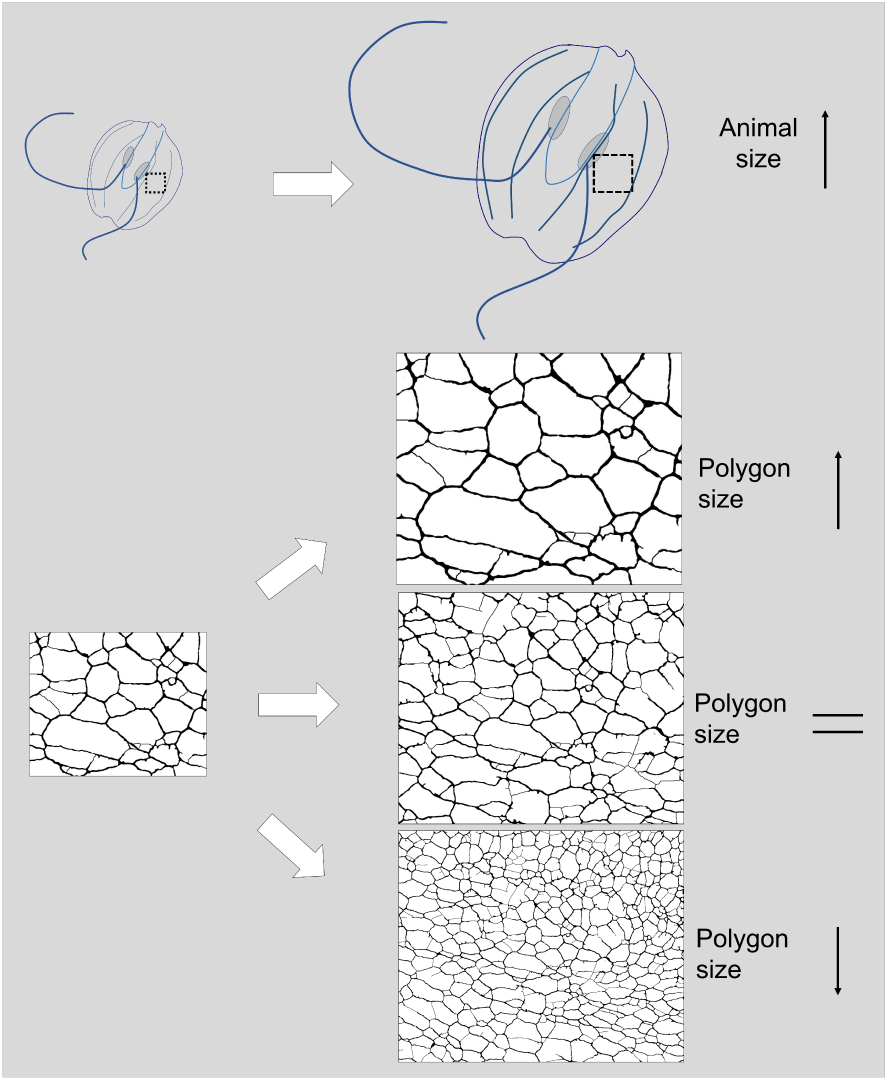
How Does the Nerve Net in *P. pileus* Restructure During Growth? This is a schematic representation of the potential effect growth will have on nerve net structure. As the animal grows the network will restructure to compensate for this change in body size. Three scenarios could be observed from the morphological characteristics of the polygons in the nerve net. The polygons refer to the area between the neural branches. The polygons could get bigger, get smaller or stay the same size. Each scenario would result from different mechanisms at the level of the individual neuron, including neurite elongation, neurite branching and neurogenesis.

To address the question of regional structural specialization in the nerve net, we needed to carry out a detailed, multi-factor characterization of the geometric and morphological characteristics of the nerve net in specific and identifiable body regions. While significant variability would be expected to exist within these regions, comparing populations of geometric descriptors between regions may reveal previously unidentified differences between those regions. Taken together, both analyses presented here represent the most detailed characterisation of ctenophore nerve net geometry and structure to date.

## Methods

### Animals

Animals used in this study were caught in Howth Harbour (53°23’34.9”N 6°03’58.0”W) and were kept in our aquaculture system (as described in Courtney et al., (2020a)) for < 24 hours before being used for this study. A total of 23 *P. pileus* were used for nerve net imaging, and an additional 6 animals were used in the validation of the body surface area estimates.

### Estimating *Pleurobrachia pileus* Surface Area

To understand the relationship between animal surface area and nerve net structure in *P. pileus*, we must first establish an accurate estimate of surface area. Direct 3D imaging techniques, such as optical projection tomography (OPT), are ideal for this task, but are not compatible with the immunofluorescent approach we used to image the nerve net. As a result, the approach we used was to model the surface area of the animals used in our nerve net analysis and use a separate cohort of animals to validate these models with whole animal 3D imaging techniques. For this we used fluorescent OPT to measure surface area of fixed animals. However, as the fixation of whole animals was likely to result in changes in body morphology and OPT in live animals was difficult to do reliably, we assessed the relationship between live and fixed animal morphology using light sheet morphometry.

Live animals were removed from tanks and their wet weight was acquired before placing on a flat surface and making axial measurements (Supplemental Figure 1B). These measurements were used to generate surface area estimates by modelling the body as a spheroid, hemisphere or partial sphere (Supplemental Figure 1C). Animals were stained with a 1:4000 solution of acridine orange (AO) in natural seawater for 10 minutes. Light sheet images were collected using a simple 0.8mm thick horizontal laser light sheet generated by placing a 7mm focal length plano-convex cylindrical lens (custom fabricated from sylgard (DOWSIL 184)) at the output aperture of an 80mw 532nm laser diode. The light sheet was directed to an acrylic tank containing seawater in which *Pleurobrachia* could swim freely. A Nikon D5600 dSLR with 50mm 1.8 lens was mounted on a tripod above the tank, and manually focused so that the plane of focus matched the plane of the light sheet. We translated the animals through the light sheet/focal plane using a motorised translating Z stage (speed: 2mm/s). Optical sections were sampled at intervals of 0.4mm and the perimeter of the animal in each optical slice was delineated manually in ImageJ (Rueden et al., 2017). The length of this perimeter was multiplied by the sampling interval (0.4mm) to estimate the surface area of that slice. For the top and bottom slices, the area was calculated directly. The calculated areas from the optical section through the animal were summed to estimate the total surface area of the live animals.

Animals were then fixed in paraformaldehyde as described by Courtney et al. (2020b). Fixed animals were again weighed, re-stained with AO, embedded in 1% agarose (Sigma-Aldrich A0169) made with distilled H_2_0 and imaged using a custom built OPT device. A telecentric lens (Thorlabs, MVTC23024) with a diagonal FOV of 45.2mm, a depth of field of ±11mm and a working distance of 103mm was fitted with a 560nm long pass filter. Fluorescent excitation was provided by a 470nm LED (ThorLabs, M470L3) collimated by aspheric condenser lenses (ThorLabs, ACL2520U-DG6-A). The agarose block containing a fixed animal was attached to a stepper motor (RS Components, 440-420) and aligned to the light path with an XYZ translating stage and suspended in distilled H_2_0. 200 RGB images at 1.8° intervals were acquired for each animal with a colour camera (FLIR, CM3-U3-13Y3C), which provided a 360° view of the animal. RGB images were converted to single channel by summing the red and green channels and inverted before reconstruction using NRecon software. Volviewer software (Avondo, 2013) was used to visualise the 3D reconstructions (Supplemental Figure 1A). The raw output of the NRecon tomographic reconstruction is a Z-slice. As the total number of Z-slices in a stack corresponded to the Z-height of the sample in pixels (and therefore 700 for the largest animals), we estimated surface area using every fourth slice to improve efficiency of the analysis. The animal boundary was outlined manually in ImageJ, the perimeter calculated and multiplied by the height (0.074mm) to calculate the surface are of that slice, and the surface areas of all slices were added to the surface of the top and bottom slices to give an estimate of the animal surface area.

### Nerve Net Imaging

For nerve net analysis, axial measurements of animals were taken prior to fixation for the purposes of surface area estimation. The tissue fixation, dehydration and immunostaining techniques were performed as described in Courtney et al., (2020b). The tissue was dissected into two preparations; the aboral and the oral dissection. The dissection procedure is outlined in Supplementary Figure 3. After dissection, tissue was permeabilized with TritonX-100 (Sigma, Aldrich, T9284) at 0.2% and 0.1% in PBS for 30 min each and immunofluorescent labelling of neurons was carried out with a fluorescent conjugated primary antibody against tyrosylated *α*-tubulin (Novus Biologicals, YL1/2, DyLight 488, NB600-506G, 1:1000 dilution). Stitched composite images of large areas of the tissue surface were then generated using a custom-built, automated eipfluorescent microscope as previously described (Courtney, Alvey, et al., 2020b).

*P. pileus* have 8 body wall planes due to their biradial symmetry, but three sampling planes (tentacular, intermediate and sagittal) are sufficient to represent all 8 planes (Figure 2A). Specific and identifiable subregions within these planes, defined by their relationship to identifiable anatomical landmarks, were then selected for analysis (Figure 2B).

**Figure 2.**
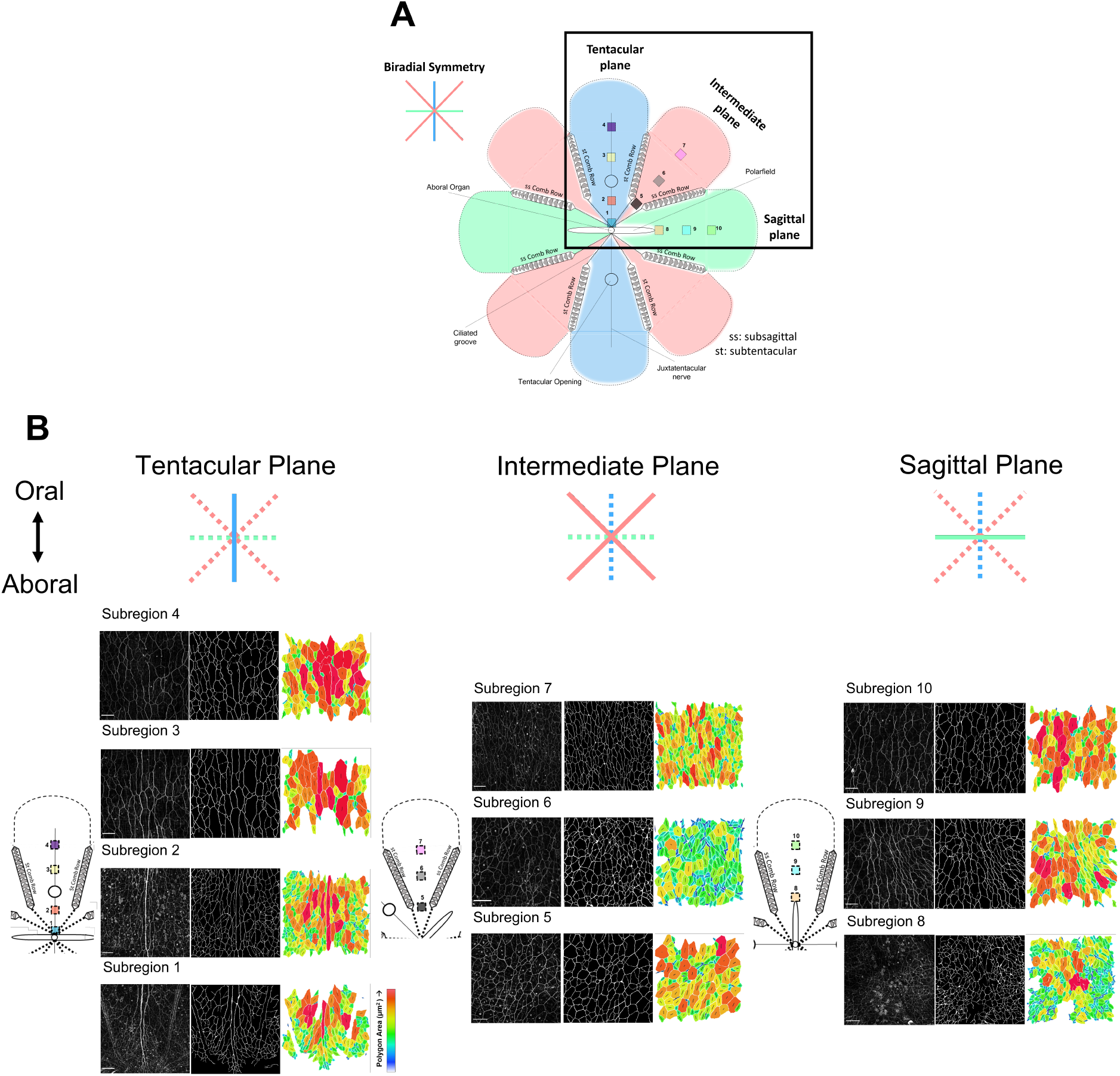
Characterising Nerve Net Subregions. (A) This is a schematic representation of *P. pileus* outer body surface including major anatomical landmarks. It is challenging to represent a spheroidal structure in 2D therefore we have positioned the aboral organ centrally and included breaks in the tissue from the oral region to the oral most comb plates. The animals have 8 body wall planes. As the animals are biradially symmetrical sampling three planes is representative of all 8 planes. Thus, the tentacular plane, the intermediate plane and the sagittal plane were denoted as the three sampling planes. Within the tentacular plane we had subregion 1-4, within the intermediate plane we had subregion 5-7 and within the sagittal plane we had subregion 8-10. With the exception of subregion 1, which is unique to the tentacular plane, subregions 2,5,8 and 3,6,9 and 4,7,10 are comparable across the planes (equivalent subregions). (B) Examples of analysed regions from all the subregions alongside the resulting segmented image. All images are orientated vertically, with the aboral region below and oral region above. To enable the visualisation of polygon morphology, size and orientation, and to maintain the spatial distribution information, we generated this third image using Matlab. This plot shows the morphology of the polygons while encoding polygon area as colour. It also includes a black line to denote orientation of a polygon. The line segment is the major axis of the polygon, while the line length is proportional to the circularity of the polygon (a shorter line denotes a more circular polygon as they possess less orientation information). Scale bar: 100 *µm*.

Within the tentacular plane subregion 1 was located adjacent to the aboral organ at the point at which the JTN vanishes into the nerve net. Subregion 2 is in between the aboral most comb plates in the tentacular plane. Subregion 3 is in between comb plates in the tentacular plane midpoint from the aboral to the oral region. Subregion 4 is in between the oral most comb plates in the tentacular plane. Within the intermediate plane, subregion 5 is in between the aboral most comb plates. Subregion 6 is in between comb plates in the intermediate plane midpoint from the aboral to the oral region. Subregion 7 is in between the oral most comb plates in the intermediate plane. Within the sagittal plane, subregion 8 is adjacent the tip of the polar field. Subregion 9 is in between comb plates in the sagittal plane midpoint from the aboral to the oral region. Subregion 10 is in between the oral most comb plates in the sagittal plane.

While the subregion locations were defined in relation to anatomical landmarks, the subregion size in each animal was defined in proportion to the animal surface area. This means that the subregion represents a specific fraction of the total surface area, rather than a specific area of absolute size, making subregions spatially comparable between animals of different sizes. A minimum region size of 800×800 pixels (308×308 *µm*) was defined for the smallest animal in the population (which had an estimated surface area of 80.56 *mm*^2^), and regions sizes were scaled proportionately for larger animals while remaining centred on the defined anatomical locations.

This analysis was carried out on a total of 23 animals, with surface areas ranging from 80.56*mm*^2^ to 517.15*mm*^2^. In each animal, any subregions that showed signs of damage or folding of the tissue which would impact the analysis were rejected. As a result, not all subregions were analysed from each animal.

### Analysis of Geometric Features of the Nerve Net

A semi-automated segmentation system was developed to isolate and define the nerve net architecture in the acquired images using Matlab. This segmentation method begins by determining optimal contrast enhancement for each image using a genetic algorithm based on an approached developed by Hashemi et al. (2010). While this algorithm is generally reliable, in some cases over and under segmentation was evident, and all images went through a process of manual checking and refinement as necessary. This segmentation process describes the nerve net as polygons with boundaries defined by the neurite bundles. As the images were acquired as maximum intensity z-projections of flat wholemount preparations we treated the network as a 2D planar structure.

#### Polygon Features

We used the following features to characterise the morphology of each polygon: area, circularity, orientation and number of polygon neighbours. The circularity of a polygon is calculated from the length of the major and minor axes, where 0 denotes a circle and 1 represents a line segment. Polygon orientation is calculated as the angle between the major axis of the polygon and the oral-aboral-axis. The number of polygon neighbours can also be interpreted as the number of sides a polygon has and thus is a measure of shape complexity.

#### Network Features

The nerve net has a branched morphology therefore we also extracted the length of each branch. This was calculated from a skeleton image of the network therefore any variability in branch thickness was disregarded. Branch length is the distance of a branch between two nodes. A node is the point at which a branch bifurcates into two daughter branches. The branching angles at all nodes were also calculated. Each node had three branches therefore the sum of the three branch angles summed to 360°.

#### Spatial Features

The above measurements do not take into account the distribution of these characteristics across the analysed region. Each analysed region was divided into an 8×8 grid and therefore constituted 64 ‘windows’. The first window was located at the top left of the analysed region, it moved to the right 8 times and then moved down to start again on the left side of the analysed region. This sequence was repeated across the entire region. Within each window we calculated the total branch length and the total number of nodes present. Using this method, the nerve net in each subregion is described by a population of large number of descriptors. To facilitate comparison between regions, we combined the descriptors to generate a feature vector for the entire nerve net population and also for each subregion. This was inspired by Farhoodi & Kording (2018) who used this approach to compare the complex 3D morphology of real and simulated neurons. The premise is to define specific morphological features and then represent the distribution of each of them as a histogram. All the distributions of all morphological features can then be combined as a feature vector, represented as a single, compound histogram. We incorporated 8 morphological features into the feature vectors; polygon area, polygon circularity, polygon orientation, polygon number of neighbours, branch length, distribution of total branch length, branching angle at nodes and distribution of nodes (Supplemental Figure 9).

As we could generate a feature vector for the entire population as well as each subregion, the degree to which each subregion differed from the overall population can be measured by generating a difference vector by subtracting the region feature vector from the population feature vector. Combining the absolute size of the components of the difference vector allowed a single estimate of overall difference between the subregion nerve net and the overall nerve net, a deviation from population (DFP) score, to be calculated. This determines which subregions are the most different from the overall population, thereby suggesting which subregions may display regional specialization.

### Data Visualisation & Statistical Analysis

The statistical analysis and data manipulation was performed using R (Core Team, 2018) and Matlab. R packages utilised for data visualisation include ggplot2 (Wickham, 2016), ggthemes (Arnold, 2019), and cowplot (Wilke, 2019).

## Results

### Does Axial Modelling Predictably Estimate Surface Area of Live *P. pileus* when Compared to OPT?

We assessed the performance of three axial models to the OPT results using linear regression (slope, p) (Supplemental Figure 1, G-I). Ellipsoid modelling generated a surface area very close to the OPT result while linear regression revealed a strong linear relationship between the methods (1.04, <0.0001). Modelling the body as a partial sphere resulted in an underestimation of surface area but it was predictable (0.59, <0.0001). The hemisphere model also performed well (0.82, <0.0001). This suggests that all three models are acceptable approaches for estimating body surface area in *P. pileus*, but correction factors would be required for the partial sphere and hemisphere models. The live ellipsoid model was used to estimate body size in animals used for nerve net analysis.

### How Does Fixation Impact Body Size and Shape?

While the live ellipsoid approach was predictably estimating surface area in a fixed animal, as nerve net imaging was carried out in fixed animals it is important to account for changes in body surface area caused by fixation. To this end, we compared the surface area measurements made using light sheet imaging in live animals to the OPT surface area measurements made in fixed animals, and found a consistently higher surface area measurement in live animals (1.23, <0.0001, Supplemental Figure 2 C,D). The fixed animals were not simply smaller, but showed evidence of shape changes, specifically prominent bulging of the body wall between comb rows (Supplemental Figure 2B), which appears to be an enhancement of a feature seen in the live animals (Supplemental Figure 2A). This artefact of fixation is likely a result of muscular shortening during fixation. We have identified that while the live ellipsoid model is predictably estimating size of a fixed animal it does not measure changes in shape.

### How Does the Nerve Net Restructure with a Change in Body Size?

The map of analysed subregions, and examples of the polygon data sampled in those regions, is shown in Figure 2B. We performed median regression analyses separately in each of the 10 subregions to examine the relationship between polygon size and animal body size. Most regions showed no statistically significant changes in polygon area with a change in body size (Figure 3). While some subregions showed evidence of small changes in size, these changes weren’t consistent in direction, ranging from decreased polygons in subregion 6 (slope= -2.78, p = 0.02) to increased polygons in subregion 9 (2.23, p<0.001). In the absence of clear, consistent and reliable changes across the entire nerve with increasing size, we cannot conclude that polygon size scales with animal size.

**Figure 3.**
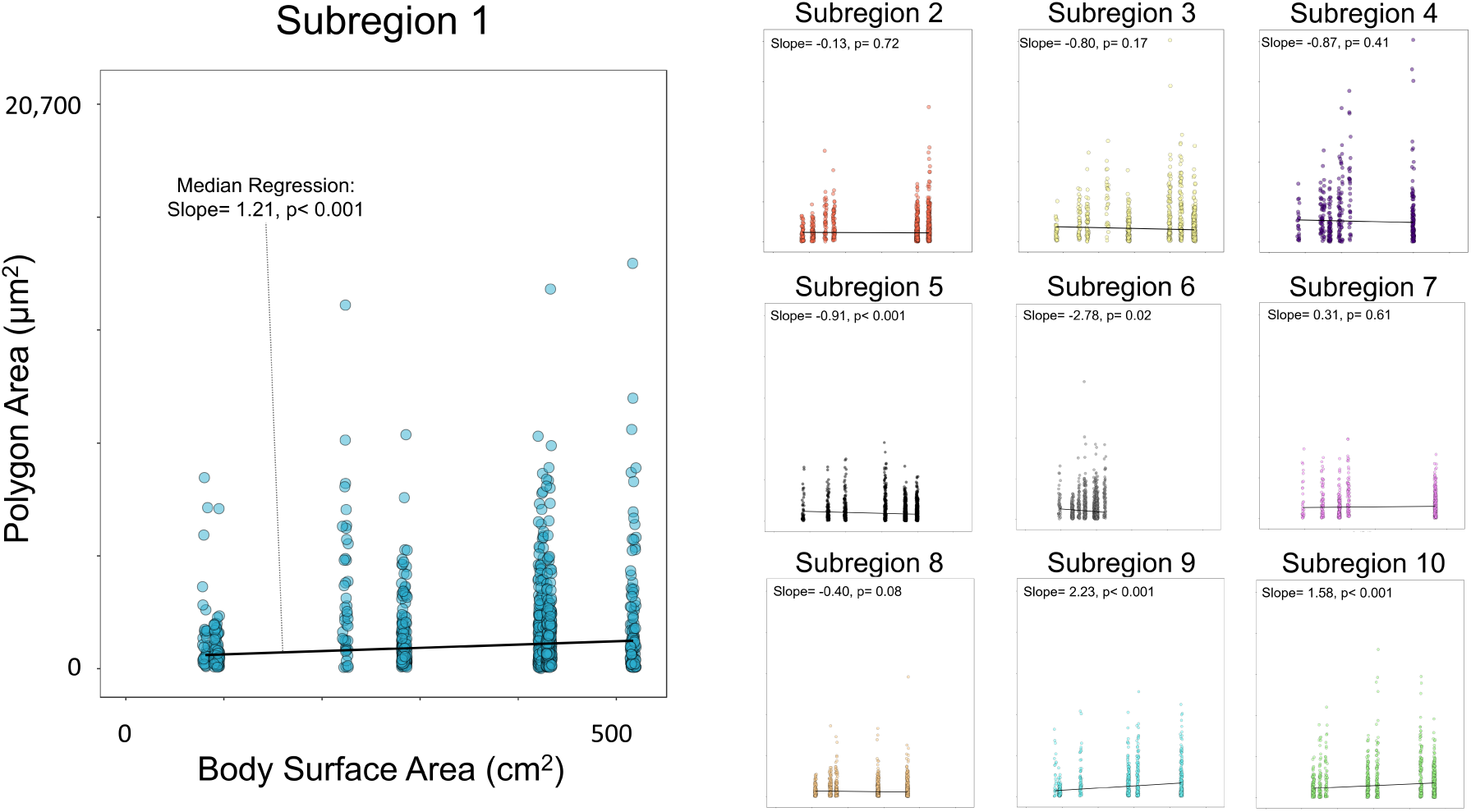
The Relationship Between Polygon Area and Body Surface Area. The relationship between polygon area and body surface area was examined by plotting polygon area (*µm*^2^) and body surface area (*mm*^2^). Body surface area was estimated using axial measurements and modelling the shape as an ellipsoid. The relationship was investigated using median regression at each subregion (black line). The slope and p-value are transcribed on each plot. A positive slope denotes an increase in polygon size with a larger body size while a negative slope would reveal the polygons getting smaller in larger animals. No clear and consistent relationships were observed between polygon area and body size across the subregions. The axes labelling conventions and limits as seen for subregion 1 are the same for all subregions.

We carried out the same analysis on polygon circularity (Supplementary Figure 4), polygon orientation (Supplementary Figure 5) and the number of polygon neighbours (Supplementary Figure 6). Again, we found no consistent differences in the shape characteristics of the polygons across animals of different sizes at various subregions across the body, suggesting no clear pattern of change in polygon shape is occurring as the animal grows. Network characteristics were also investigated in the context of restructuring with a change in body size. Both branch length (Supplementary Figure 7) and branching angles (Supplementary Figure 8) at nodes were examined. For branch length, most of the results were not statistically significant and the rest of the changes were small. For branching angles, none of the models were statistically significant. This implies that the geometric properties of the neurites within the network are unchanged in response to growth.

### How Does the Nerve Net Morphology Vary at Different Parts of the Body?

#### Comparing the Distributions of the Morphological Features in Isolation

The architecture of the network appears to be remarkably consistent across animals of different sizes at all subregions sampled. However, it is unclear if there are quantifiable differences in morphological features between subregions. Previous accounts of ctenophore nerve nets alluded to some regional specialisations in terms of structure (Jager et al., 2011), such as the presence of the JTN. This feature suggests that information is flowing longitudinally in the aboral-oral plane, but directionality is unclear. The intermediate and sagittal planes do not possess a nerve-like structure, but do they possess other features which imply signal propagation in this orientation? Jager et al., (2011) also noted that polygon density and orientation was variable at different parts of the body. We wanted to compare differences in morphology *within* each plane and *between* planes in ‘equivalent’ subregions (Figure 4).

**Figure 4.**
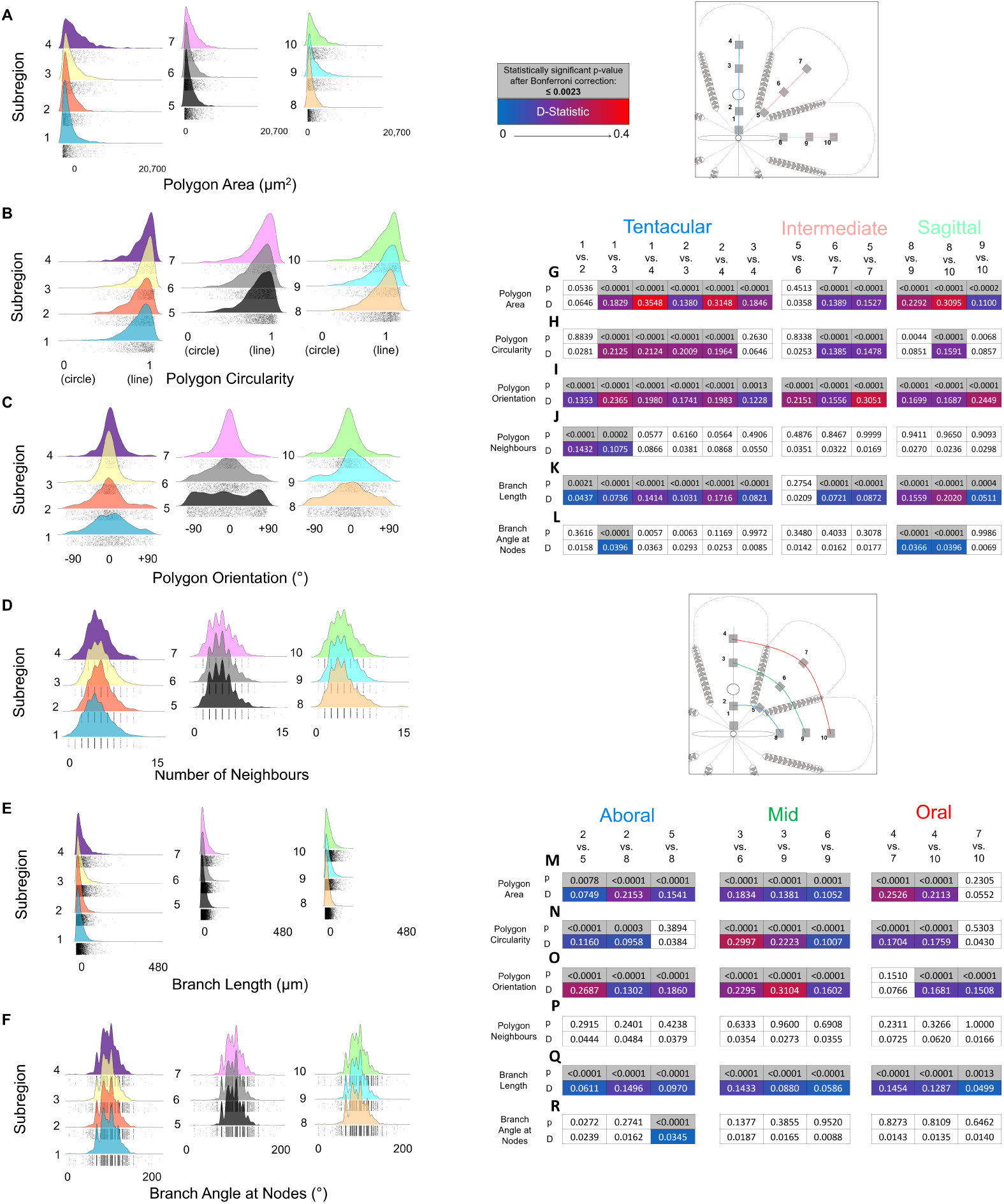
Analysing the Individual Frequency Distributions of Each Nerve Net Morphological Feature Within a Subregion and Between Subregions. Is the nerve net architecture variable at different parts of the body? (A) The frequency distribution of polygon area by subregion. (B) The frequency distribution of polygon circularity by subregion. (C) The frequency distribution of polygon orientation by subregion. (D) The frequency distribution of number of polygon neighbours by subregion. (E) The frequency distribution of branch length by subregion. (F) The frequency distribution of branch angle at nodes by subregion. The individual data points are seen along the bottom of each distribution. We performed K-S tests on all subregions within a plane (G-L) and between planes (M-R) to assess the variability of the 6 nerve net morphological characteristics. The p-value was Bonferroni corrected and the critical threshold to reject the null hypothesis was set at 0.0023. The D-statistic is a scale from 0-1 and equates how different the distributions are. Statistically significant p-values can be seen in grey and the D-statistic is colour coded from 0 (blue) to 0.4 (red).

To this end, we analysed the frequency distributions of all 6 nerve net morphological features separately for each subregion. We performed Kolmogorov-Smirnov (K-S) tests for all subregions within a plane and between planes. As this analysis involved multiple pairwise comparison, we controlled for the familywise error rate by adjusting the alpha using a Bonferroni correction. Subregions 1-4 resides in the tentacular plane and this results in 6 K-S tests per morphological feature comparing each subregion to all the others. Subregions 5-7 are in the intermediate plane and this equates to 3 K-S tests. Subregions 8-10 are in the sagittal plane and this equates to 3 K-S tests. Subregions 2,5,8 are equivalent in all planes and near the aboral region of the body, 3 K-S tests were performed to compare each subregion in this group to the other. Subregions 3,6,9 are equivalent subregions mid-way from the aboral to the oral part of the body. Subregions 4,7,10 are equivalent subregions near the mouth. The total number of comparisons in the family is therefore 21, and so alpha was adjusted to 0.0023.

The frequency distribution for polygon area is skewed with most polygons falling within a size range of ∼50-1600*µm*^2^ (Figure 4A) but larger polygons are observed on occasion (up to 20,916*µm*^2^). K-S tests of all the subregions within a plane showed statistically significant differences in the distributions (Figure 4G). The highest D-statistics for all the morphological features across all subregions was observed for polygon area. When comparing between planes polygon area displayed statistically significant differences in all comparisons except 7/10 (Figure 4M). This suggests that most subregions have distinct and specific profiles of polygon sizes.

The frequency distribution for polygon circularity is skewed indicating that most of the polygons are more elongated than circular (Figure 4B). K-S tests for most of the comparisons within a plane (Figure 4H) and between planes (Figure 4N) showed statistically significant differences in the distributions. Polygon circularity appears to be regionally specific to some extent.

The frequency distribution shape for polygon orientation across all subregions indicates an overall tendency to align to the oral-aboral axis (Figure 4C). The distributions of 1,2,5 and 8 are flatter and represent the orientation of the polygons fanning left and right of the oral-aboral-axis in these areas. Subregions 3,4,6,7,9 and 10 have narrower distributions indicating that more polygons are oriented in the oral-aboral-axis. This provides structural evidence that the network is aligned in aboral-oral plane and may suggest information flowing longitudinally in the body wall between the comb rows in all three planes. The K-S tests statistically analysed the distribution within a plane and found that they were all different (Figure 4I). When comparing between planes all comparisons (except 4/7) had different distributions (Figure 4O). A directional bias which is variable across body regions appears to be an organisational principle of the nerve net.

The number of polygon neighbours (or number of sides) is most often between 4 and 7 but up to 16 neighbours can occur in rare instances (Figure 4D). The K-S test revealed statistically significant difference in 1/2 and 1/3 but no other comparisons within (Figure 4J) or between planes (Figure 4P) were different. The number of polygon neighbours is a proxy for number of sides on a polygon and in turn is a measure of shape complexity. This feature appears to be largely conserved across the subregions.

Branch length distribution is skewed with most branches falling within the 1-32*µm* size range, but some long branches (up to 444*µm*) do occur (Figure 4E). The K-S tests revealed that all comparisons within planes (Figure 4K, except 5/6) and between planes (Figure 4Q) had statistically significant differences in their distributions. The D-statistics were generally low and therefore the differences between regions is small.

The frequency distributions of branch angles at nodes have a small peak at 90° but generally fall within 100° and 130° (Figure 4F). The K-S tests showed that only three of the comparisons within planes had statistically significant differences in their distributions and the effect was small (Figure 4L). When comparing between planes only one difference (5/8) was noted (Figure 4R). It appears that a relatively consistent mode of bifurcation is occurring at nodes across the entire network.

All 6 nerve net morphological features displayed some degree of variability at different parts of the body. While some features were more variable than others, generally the effects were subtle and small. However, the features are not independent of each other and some are intrinsically linked. Studying each feature in isolation may not be the best way to compare a complex repertoire of morphological characteristics. Assessing the impact of all features simultaneously would be preferable. To this end, we generated a feature vector which combines all morphological features for each subregion.

#### Feature Vectors

The feature vector includes 8 morphological features; the 6 features we have been referencing until this point and two additional features, spatial distribution of nodes and branches, which describe the uniformity of the network by dividing the image into an 8×8 grid and measuring the node number and total branch length in each region of the grid. A feature vector is a combination of histograms for each feature (Supplemental Figure 9). Each histogram has a specific number of bins, a bin width and an upper/lower bin limit. All the feature histograms were combined to make a feature vector for each subregion (Figure 5). The feature vectors demonstrated differences in polygon area, particularly in subregion 4 which possessed more large polygons than other subregions. Some subregions displayed a relatively constant frequency in the spatial distribution features while others showed some patterns of lower frequencies at the lateral parts of the analysed region.

**Figure 5.**
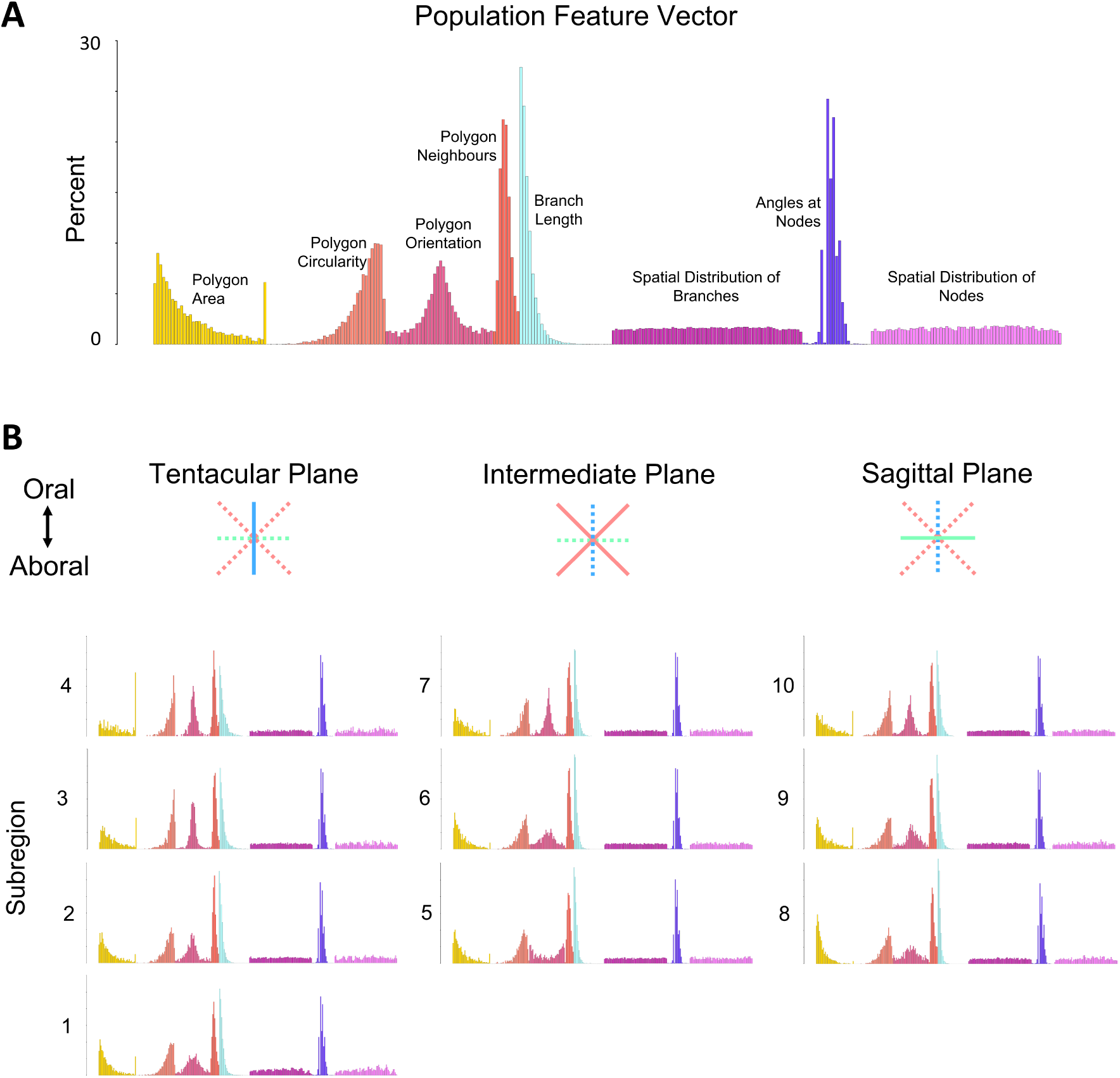
Population and Subregion Feature Vectors. 8 morphological features were extracted from the analysed regions of the nerve net. They include polygon area, polygon circularity, polygon orientation, polygon number of neighbours, branch length, spatial distribution of total branch length, branching angle at nodes and spatial distribution of nodes. (A) The population feature vector was generated by summing all subregion feature vectors and expressing the densities as percent. (B) Subregion feature vectors grouped by body plane. Subregion feature vectors were generated by summing all the analysed regions that make up a subregion and expressing the densities as percent. The labelling conventions and axes limits are the same for all feature vectors.

All the subregions were combined to generate a population feature vector (Figure 5A). Each subregion was compared to the population vector to generate a difference vector (Figure 6) and a deviation from the population score. The difference vectors show a degree of variation in nerve net structure across the body, with the most pronounced differences evident in polygon size, circularity and orientation. The largest DFP score was recorded for subregion 4 (Figure 6 C,D), which is the most orally positioned region in the tentacular plane. Overall, our data reveal that differences between body regions are relatively modest and that the nerve net morphology is largely conserved across the body.

**Figure 6.**
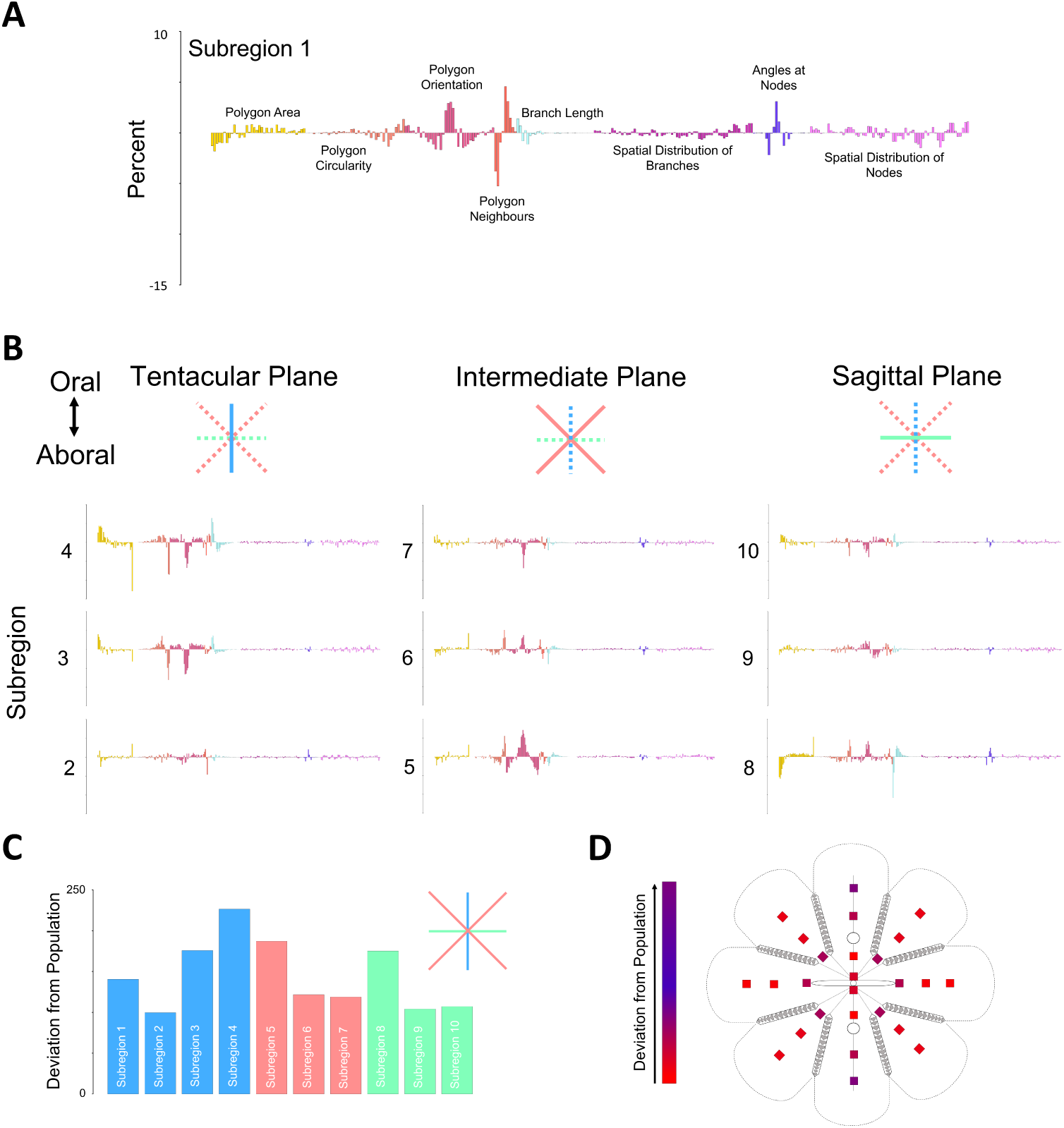
Subregion Difference Vectors. In order to examine the difference in network architecture across different parts of the body the feature vectors for the subregions and the population feature vector were compared to generate a difference vector. The population feature vector is subtracted from the subregion feature vector to generate a subregion difference vector. This difference vectors shows how variable a subregion is from the entire population. The labelling conventions and axes limits are the same for all difference vectors. (A) Subregion 1 difference vector. (B) Subregion 2-10 difference vectors grouped by body plane. Summing a subregion difference vector resulted in a deviation from population score. (C) Subregion deviation from population scores, coloured by plane. (D) A schematic representation of *P. pileus* external body surface modelled in 2D with major anatomical structures included and all 10 subregions labelled in all planes. The deviation from population score is coloured in a scale from red (lowest) to purple (highest).

### What are the Relationships Between the Nerve Net Morphological Features?

While the feature vectors provide a detailed statistical description of the complex morphology of a nerve net, they do not include all the relevant information. For example, the relationship between variables cannot be captured in a feature vector. To test whether there is a relationship between morphological features and whether those relationships varied across the subregions, we examined the relationship between polygon orientation and circularity (Figure 7), polygon area and circularity (Figure 8) and polygon area and number of neighbours (Figure 9).

**Figure 7.**
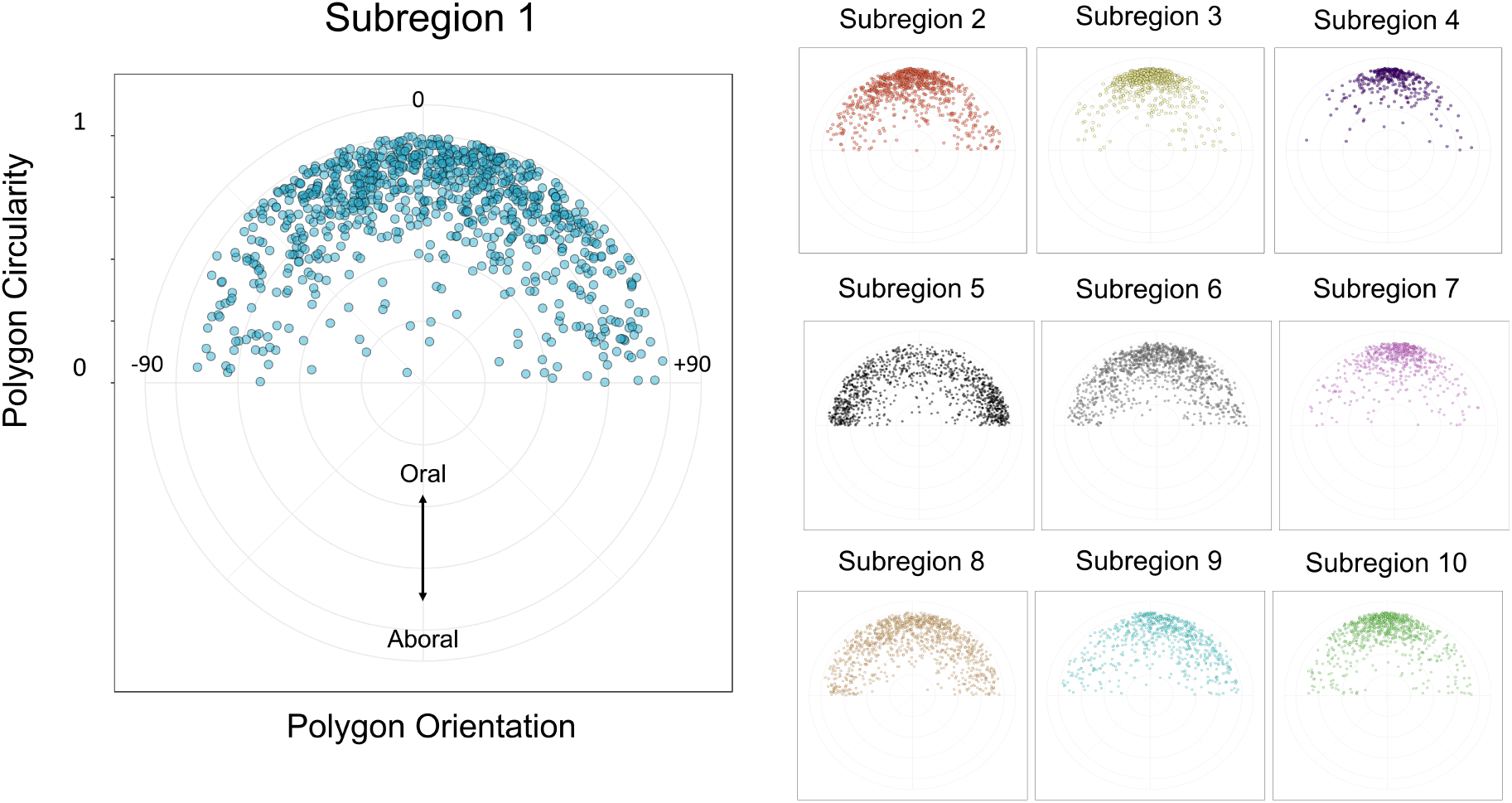
The Relationship Between Polygon Circularity and Polygon Orientation. The relationship between polygon circularity and polygon orientation was examined. The circularity of a polygon is calculated by a ratio of the minor axis to the major axis. This results in a value between 0 and 1 (0 denotes a circle while 1 represents a line segment). Polygon orientation is calculated as the angle of the major axis to the 0° axis. For all analysed regions, the 0° axis was defined as the oral-aboral-axis. The axes labelling conventions and limits as seen for subregion 1 are the same for all subregions.

**Figure 8.**
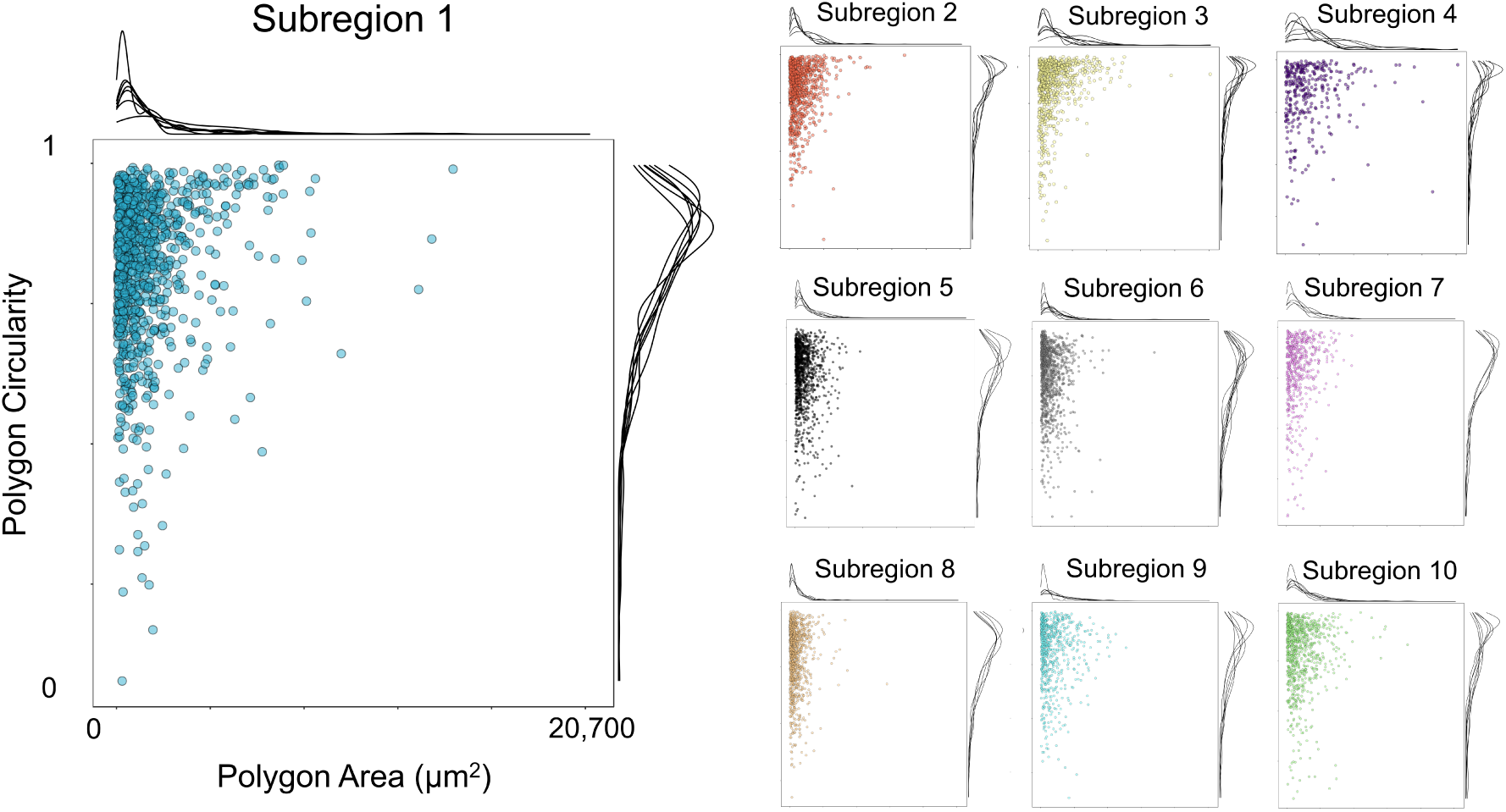
The Relationship Between Polygon Circularity and Polygon Area. The relationship between polygon circularity and polygon area was examined. The circularity of a polygon is calculated by a ratio of the minor axis to the major axis. This results in a value between 0 and 1 (0 denotes a circle while 1 represents a line segment). A frequency distribution for each individual analysed region (animal) is seen for both variable on the margins of the plots. The axes labelling conventions and limits as seen for subregion 1 are the same for all subregions. Larger polygons tend to be less circular and more oblong in shape. This finding is consistent across all subregions.

**Figure 9.**
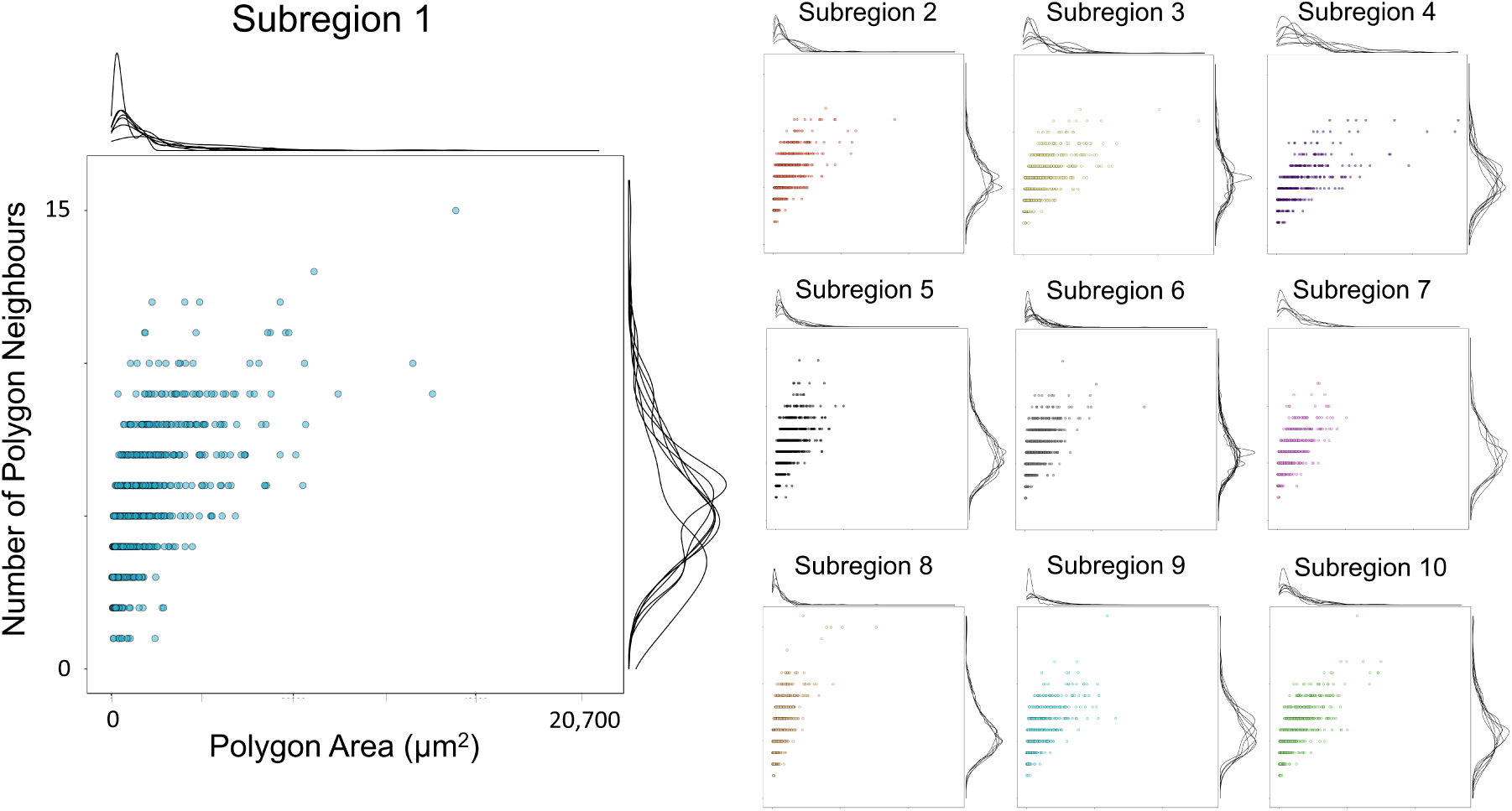
The Relationship Between Polygon Area and Polygon Neighbours. The relationship between polygon area and number of polygon neighbours was examined. The number of polygon neighbours can also be interpreted as the number of sides a polygon has and thus a measure of shape complexity. A frequency distribution for each individual analysed region (animal) is seen for each variable on the margins of the plots. The axes labelling conventions and limits as seen for subregion 1 are the same for all subregions. Larger polygons tend to have more neighbours.

The circularity of a polygon and the orientation are intrinsically linked. A polygon with a less circular and more oblong shape carries more orientation information. The polygons orientated with the oral-aboral-axis appear to be less circular and have more of an oblong shape (Figure 7). The less circular polygons also appear to be generally larger (Figure 8). In addition, larger polygons seem to be more likely to have more neighbours (Figure 9). We saw no clear evidence of any difference in any of these relationships between the subregions.

## Discussion

We have provided a comprehensive account of the nerve net morphology of *Pleurobrachia pileus* in animals of different sizes and across different parts of the body. A total of 68 regions were analysed with 23 animals used to generate 5-10 examples for each of the 10 subregions. This equated to 8680 polygons and ∼740mm total branch length. Our findings suggest that a stereotypical morphological arrangement of the nerve net is largely conserved throughout growth. We have also established the principles of organisation of the network and showed that some of the morphological features are variable across the subregions.

### *Pleurobrachia pileus* Displays Remarkable Stereotypy in the Structure of the Epithelial Nerve net in Animals of Different Sizes

We hypothesised that as the animal got larger, the neurites would increase in length, resulting in proportionately larger polygons. Instead we found that the polygons remained largely constant in size and shape despite the body surface area getting larger, which implies an increase in the total number of polygons. While we do not know exactly what process underpins this, the two alternatives are the formation of new polygons by existing neurons increasing their structural complexity, or by maintaining neuron shape and complexity while adding new neurons to the network to form the new polygons. Both phenomena have been observed in other species. This mechanism of adding neurons during growth was observed in two cnidarian species, *Hydra* and *Nematostella* (Bode et al., 1973; Havrilak et al., 2017).

However, the alternative (increasing complexity but not number of neurons) seems more common in species studied to date. This has been with growth in the majority of species studied to date. For example, *Drosophila* appear to maintain function by restructuring existing neurons to compensate for a changing body size. The addition of new neurons is relatively low in *Drosophila* larvae from hatching to pupariation (Truman & Bate, 1988) and qualitatively no major changes in behaviour are observed (Almeida-Carvalho et al., 2017). Grueber et al. (2002) found that dendritic arborizations of mechanosensory neurons in *Drosophila* larvae grow to maintain receptive fields, while Davis & Goodman (1998) observed that synaptic efficiency at neuromuscular junctions is altered to compensate for changes in muscle size during development. A comparative connectomics approach saw that neuron morphology changed during growth, but connectivity was fairly consistent (Gerhard, Andrade, Fetter, Cardona, & Schneider-Mizell, 2017). This phenomenon has also been described in the Cnidaria. Havrilak et al. (2017) examined neural subtypes within the nerve net of *Nematostella* and revealed that their number, morphology and location are minimally variable during development. Neither strategy seems strongly associated with particular phylogenetic groups, nor are the strategies mutually exclusive, so both may be operating in ctenophores.

### Behaviours During Growth and Network Restructuring

Why would an animal maintain certain fixed and stereotypical nervous system structures throughout significant body growth? Could the conservation of structure relate to the conservation of function? *Pleurobrachia* displays a relatively small repertoire of simple behaviours (Courtney, Merces, et al., 2020a), and although it has never been directly investigated, it seems likely that behaviours are largely conserved throughout the animal’s life. The feeding behaviour has been demonstrated in adult and larval forms (Anthony G. Moss, 1991). Global ciliary reversal was observed in larval forms (Tamm & Tamm, 1981) but it has not been reported in adults in the literature. However, we observed it on occasion in adults in our colony. The neural framework to control these important behaviours which pertain to eating and avoiding noxious stimuli must be laid down early on and conserved throughout life. Could morphological features like polygon size be functionally important elements of this framework? From a sensory perspective, increased branching to maintain polygons of a specific size could enable consistent coverage of a receptive field of a specific size. From a motor perspective, the neurites may extend their branches to recruit additional muscle fibres into their motor unit. Whether additional muscle fibres are added or whether the fibres get bigger in ctenophores during growth is also not understood however.

### Regional Variation in Nerve Net Morphology

When comparing subregions within and across the body planes, we observed subtle differences as well as commonalities in nerve net morphology. Certain characteristics such as polygon area, polygon circularity and branch length demonstrated differences in their distributions which suggests that the organisational principles governing these features are regionally specific. The number of polygon neighbours and branch angles at nodes appeared to be more stereotyped and have conserved rules across subregions. Identifying features that vary across body regions could suggest that different parts of the body are undertaking specialised functions. On the other hand, identifying the features that generalise allows us to infer fundamental properties of the network that must be important for general functionality, or result from critical developmental rules. Again, if the nerve net morphology is functionally important, these observations might reflect regional functional specialization within the nerve net.

### Is *Pleurobrachia pileus* a Viable Model Organism for Whole Organism Network Analysis?

The key missing information which would help answer many of the questions posed by our results is the morphology, distribution and connectivity of individual neurons in the network. The study presented here explicitly does not address this question. While we believe that *Pleurobrachia* has many properties that make it a useful model organism for neuroscience, one of the drawbacks compared to the more established model organisms is the lack of well validated experimental tools. We used immunofluorescent labelling of neurons as the best currently available technique, but it does have limitations. The antibody used for nerve net visualisation displayed characteristic non-specific binding which meant that automated segmentation was challenging, and significant manual validation was required. This manual step in the image analysis pipeline is not ideal given the time-consuming nature, and the risk of inconsistencies and potential biases affecting the data (Watters, Pickering, Murphy, Murphy, & O’Connor, 2014). We also cannot rule out the fact that some neurons may not have been stained with this method (Lin, Gallin, & Spencer, 2001). Importantly, it does not allow us to identify single cells in the network with a high degree of certainty, nor does it allow us to identify synaptic sites.

We have demonstrated that *P. pileus* is experimentally amenable in the context of neuroanatomical investigations. Our analysis may provide insights into the rules that govern the organisation of a nerve net as we identified key morphological features which are conserved during growth and across different parts of the body. We also identified structural directionality within the network and future work may be able to associate these neural pathways with specific behaviours. We hope this work also helps motivate the wider adoption of these organisms and the development of new, specific research tools which will bring us closer to understanding how a nervous system operates on a whole organism level from neuron to network to behaviour.

## Acknowledgements

This work is financially supported by School of Medicine, University College Dublin and European Research Council (ERC) under the European Union’s Horizon 2020 Research and Innovation Programme (Grant ERC-2014-CoG-646923-DBSModel). The authors also wish to acknowledge Dominic Courtney for his invaluable assistance in the collection of ctenophores.

