## Supplementary Figure for "Characterisation of Geometric Variance in the Epithelial Nerve Net of the Ctenophore *Pleurobrachia pileus*"

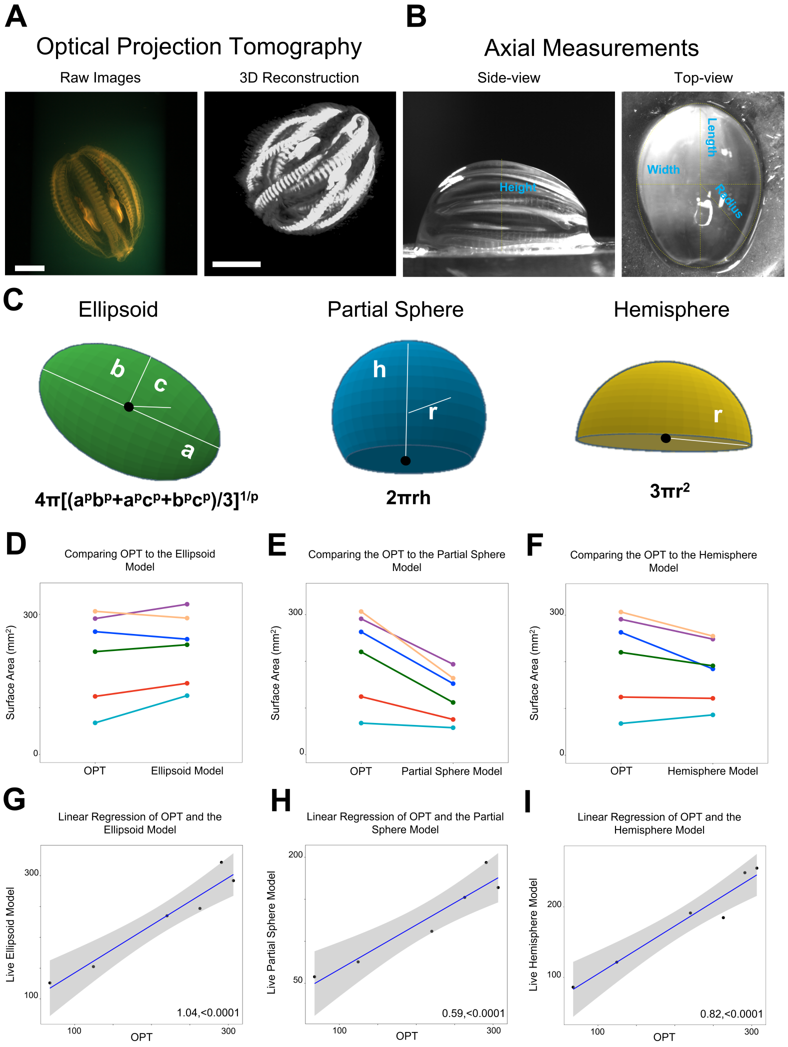


**Supplemental Figure 1. Estimating *Pleurobrachia pileus* Body Size: Comparing Axial Models to Optical Projection Tomography.** Optical Projection Tomography is the gold standard method for reconstructing mesoscales structure in 3D and is therefore the most accurate means of estimating surface area of *P. pileus*' body. Animals that were used for immunostaining experiments could not be stained with acridine orange or embedded in agarose (as was required for OPT). Therefore, we needed an accurate surface area estimation method that would not compromise the tissue quality for animals needed for nerve net analysis. (A) During OPT acquisition raw images were acquired of animal from different orientations and then they could be reconstructed in 3D. (B) We acquired images of live animals temporarily out of liquid from a top- and side-view which allowed us to extract axial measurements. (C) Axial model surface area equations. We then compared the three axial modelling methods to our OPT method to assess how predictable they are at estimating surface area. (D) Comparing the surface area estimated from OPT to the surface area estimated using the ellipsoid model from live axial measurements. (E) Comparing the surface area estimated from OPT to the surface area estimated using partial sphere model from live axial measurements. (F)Comparing the surface area estimated from OPT to the surface area estimated using the hemisphere model from live axial measurements. (G) Linear regression of the surface area estimated from OPT to the surface area estimated using the ellipsoid model from live axial measurements (H) Linear regression of the surface area estimated from OPT to the surface area estimated using partial sphere model from live axial measurements. (I) Linear regression of the surface area estimated from OPT to the surface area estimated using the hemisphere model from live axial measurements. The linear regression plots display the model as a blue line and the confidence intervals as the grey area. The slope and p-value are transcribed on each plot.

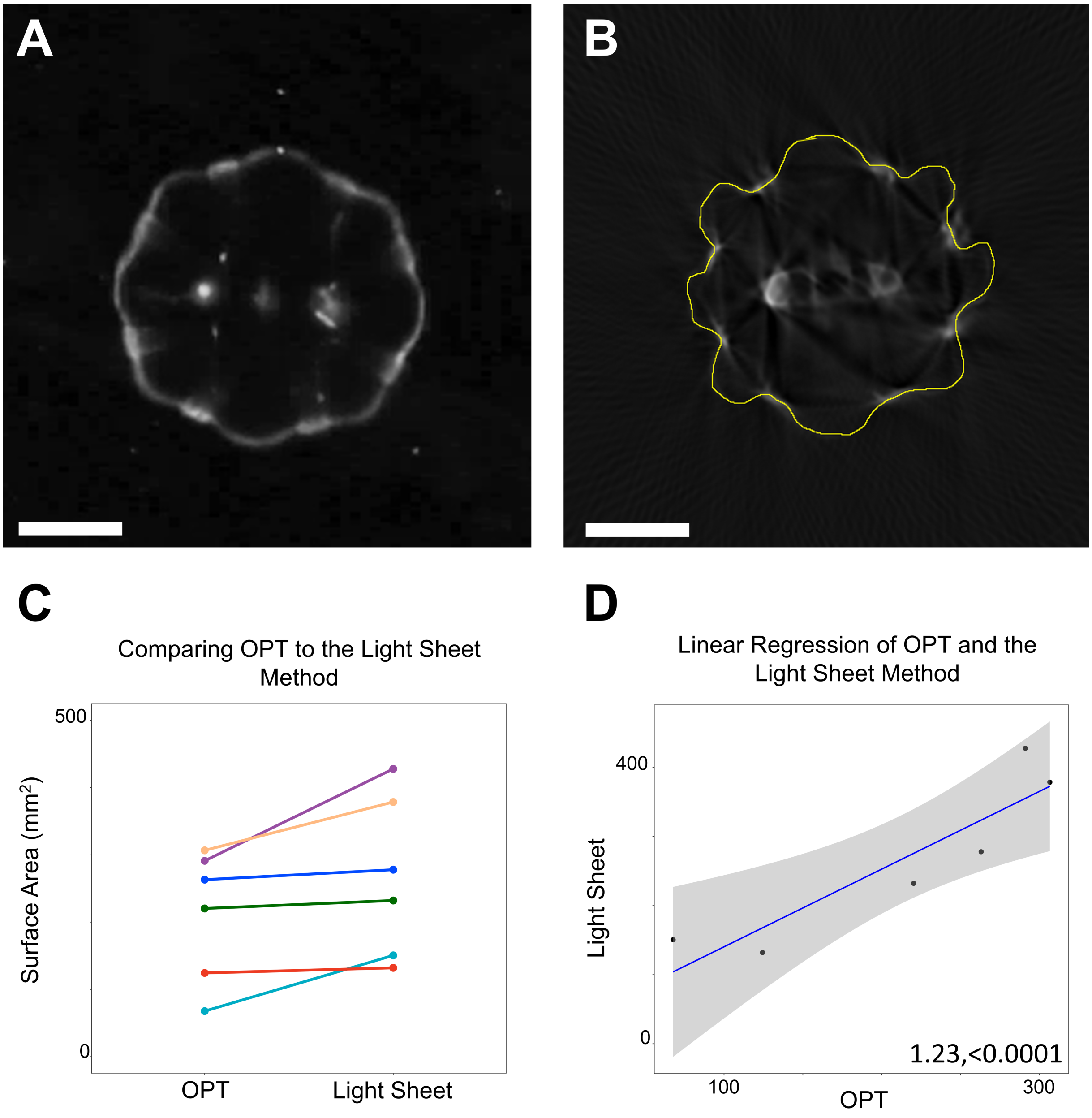


**Supplemental Figure 2. Estimating *Pleurobrachia pileus* Body Size: Comparing Light Sheet Morphometry to Optical Projection Tomography.** OPT could not be performed on the same animal in a live and fixed state therefore we built a custom light sheet morphometry system and acquired equivalent Z-slices of the animals while they were alive. We wanted to understand how fixation is affecting body shape. (A) Light sheet morphometry example Z-slice. (B) OPT Z-slice in the same animal seen in A. In between the comb rows prominent bulging can be observed when the animal is fixed but not when they are alive and thus it appears that this is a fixation artefact. (C) Comparing the live light sheet method to the fixed OPT method. (D) Linear regression of the live light sheet method and the fixed OPT method. The linear regression plots display the model as a blue line and the confidence intervals as the grey area. The slope and p-value are transcribed on each plot. Scale bar: 3mm.

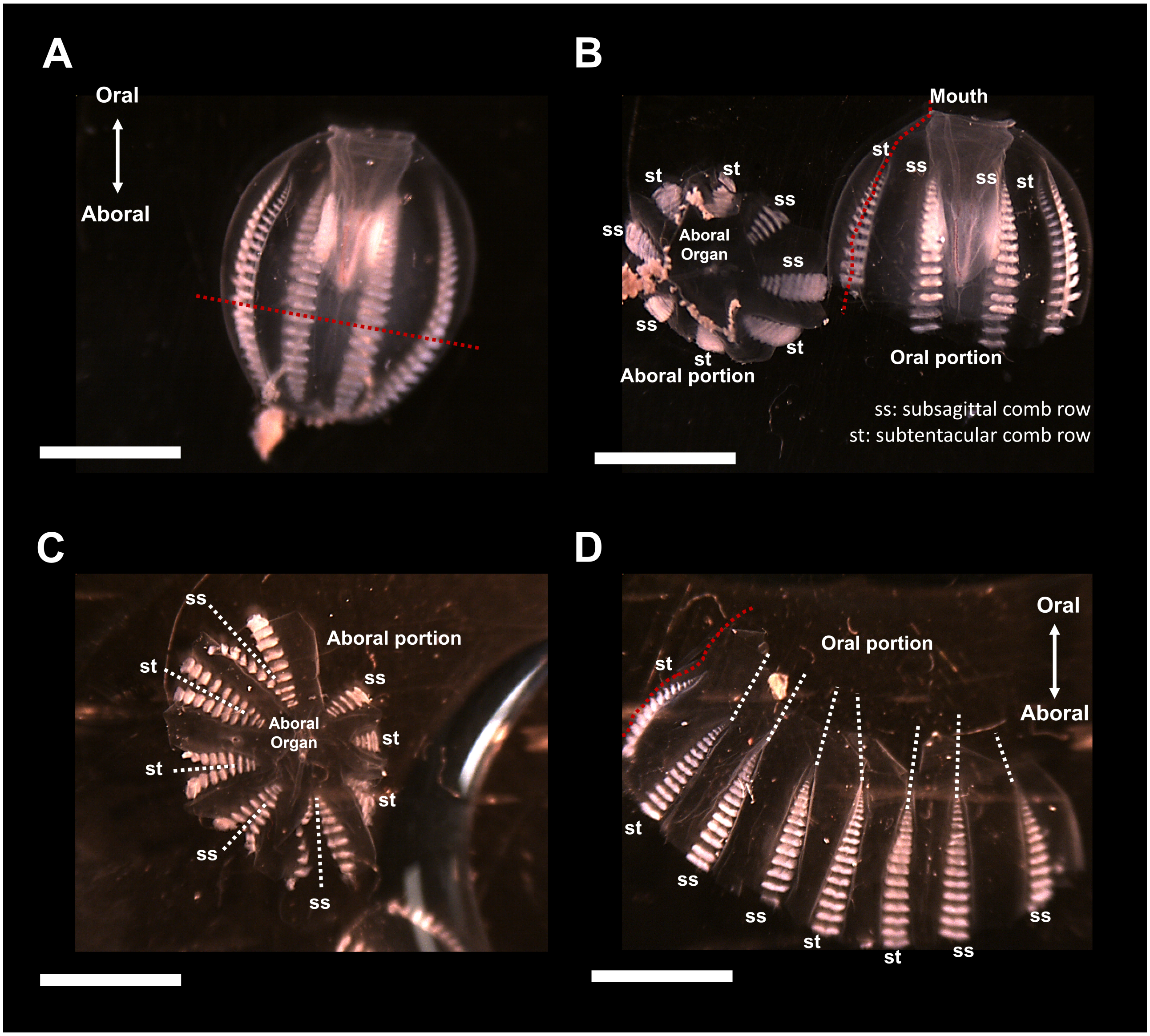


**Supplemental Figure 3. Wholemount Tissue Processing.** Wholemount tissue preparations were used and were achieved by first cutting the animal into two preparations; the aboral and the oral dissection. (A & B) The aboral dissection is prepared by cutting transversely through the entire body near the aboral most tips of the comb rows (red dashed line). (C) Cuts were made between each comb row (lateral to medial) (dashed white lines). (B & D) The oral dissection is prepared by taking the other half of the animal's body and cutting longitudinally between a ss and st comb row (red dashed line).To improve flat mounting, cuts were also made from the edge of the tissue to the tips of the oral most comb plates (white dashed lines). All internal anatomical structures were teased away to improve flat mounting of both dissected halves.

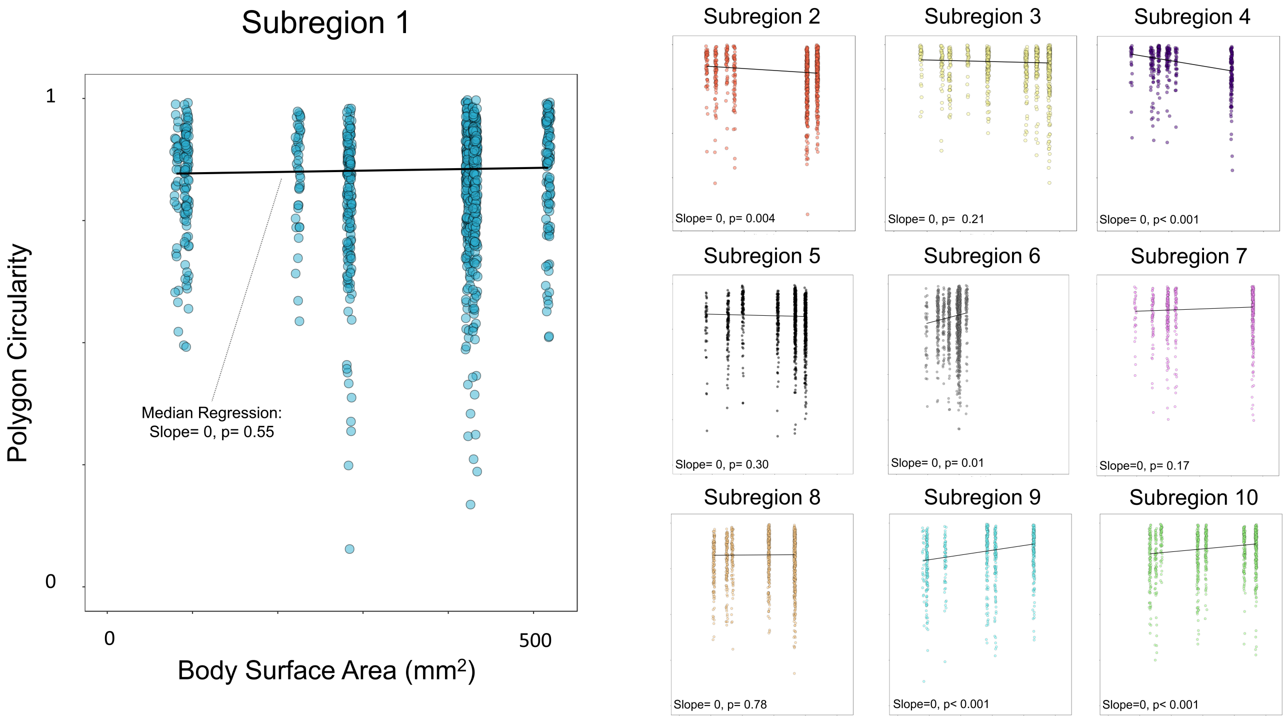


**Supplemental Figure 4. The Relationship Between Polygon Circularity and Body Surface Area.** The relationship between polygon circularity and body surface area was examined. Body surface area was estimated using axial measurements and modelling the shape as an ellipsoid. The relationship was investigated using median regression at each subregion (black line). The slope and p-value are transcribed on each plot. A positive slope denotes more oblong polygons with a larger body size while a negative slope would reveal the polygons were more circular in larger animals. No clear relationship was observed between polygon circularity and body size in any of the subregions. This suggests that the polygon circularity is constant in animals of different sizes. The axes labelling conventions and limits as seen for subregion 1 are the same for all subregions.

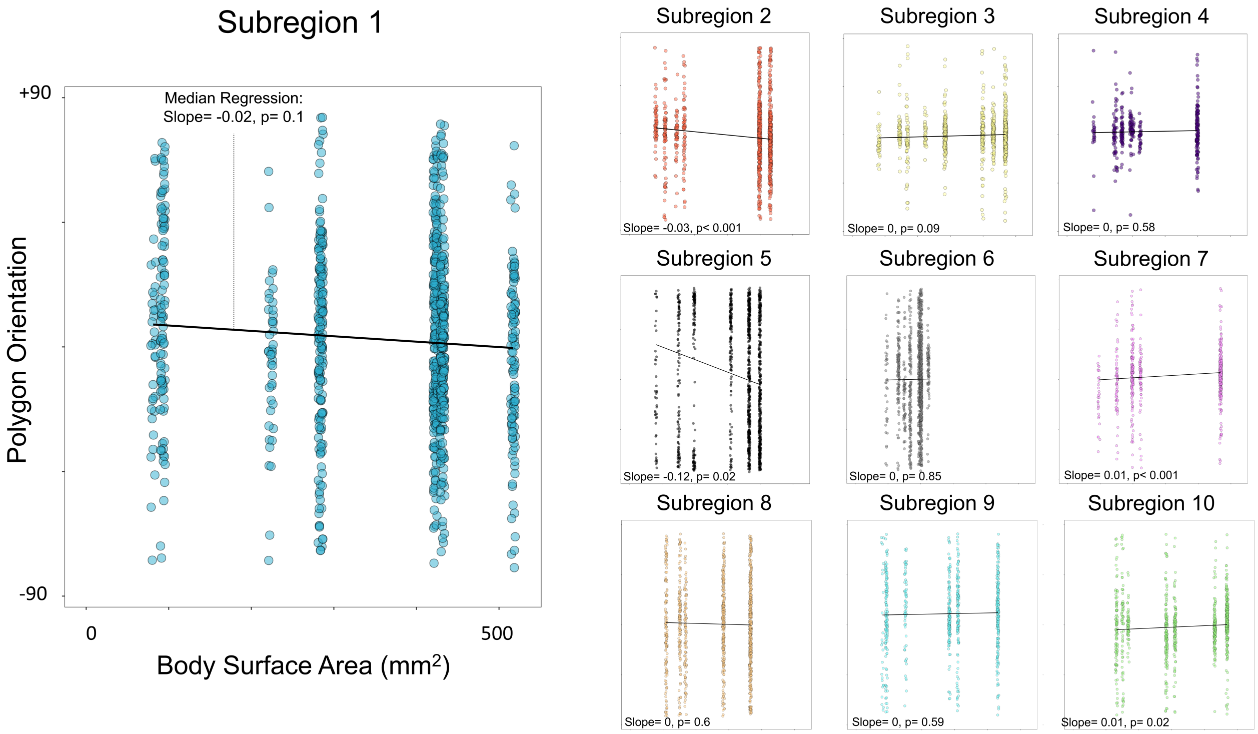


**Supplemental Figure 5. The Relationship Between Polygon Orientation and Body Surface Area.** The relationship between polygon orientation and body surface area was examined. Body surface area was estimated using axial measurements and modelling the shape as an ellipsoid. The relationship was investigated using median regression at each subregion (black line). The slope and p-value are transcribed on each plot. No relationship was observed between polygon orientation and body size in any of the subregions. This suggests that the polygon orientation is conserved across animals of different sizes. The axes labelling conventions and limits as seen for subregion 1 are the same for all subregions.

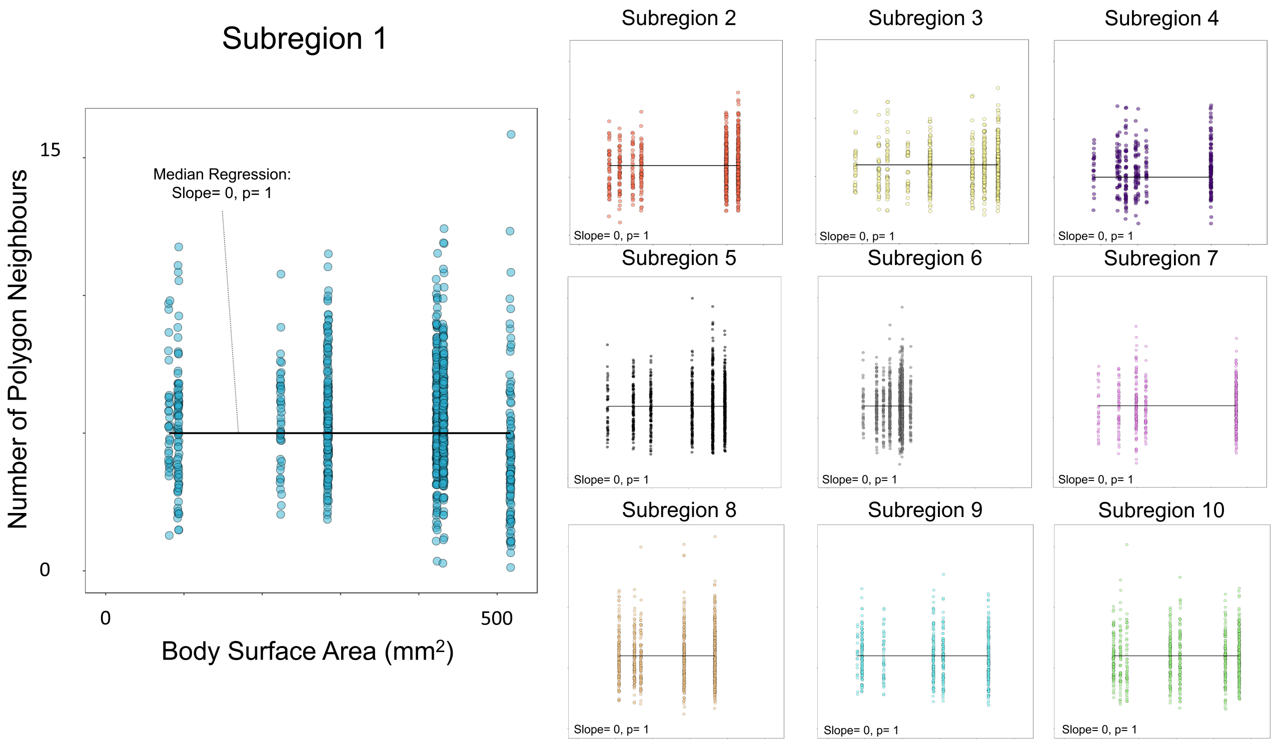


**Supplemental Figure 6. The Relationship Between Number of Polygon Neighbours and Body Surface Area.** The relationship between the number of polygon neighbours and body surface area was examined. Polygon neighbours is a proxy for number of polygon sides and is therefore also considered a measure of shape complexity. Body surface area was estimated using axial measurements and modelling the shape as an ellipsoid. The relationship was investigated using median regression at each subregion (black line). The slope and p-value are transcribed on each plot. A positive slope denotes more neighbours with a larger body size while a negative slope would reveal there to be less neighbours in smaller animals. No relationship was observed between number of polygon neighbours and body size in any of the subregions. The axes labelling conventions and limits as seen for subregion 1 are the same for all subregions.

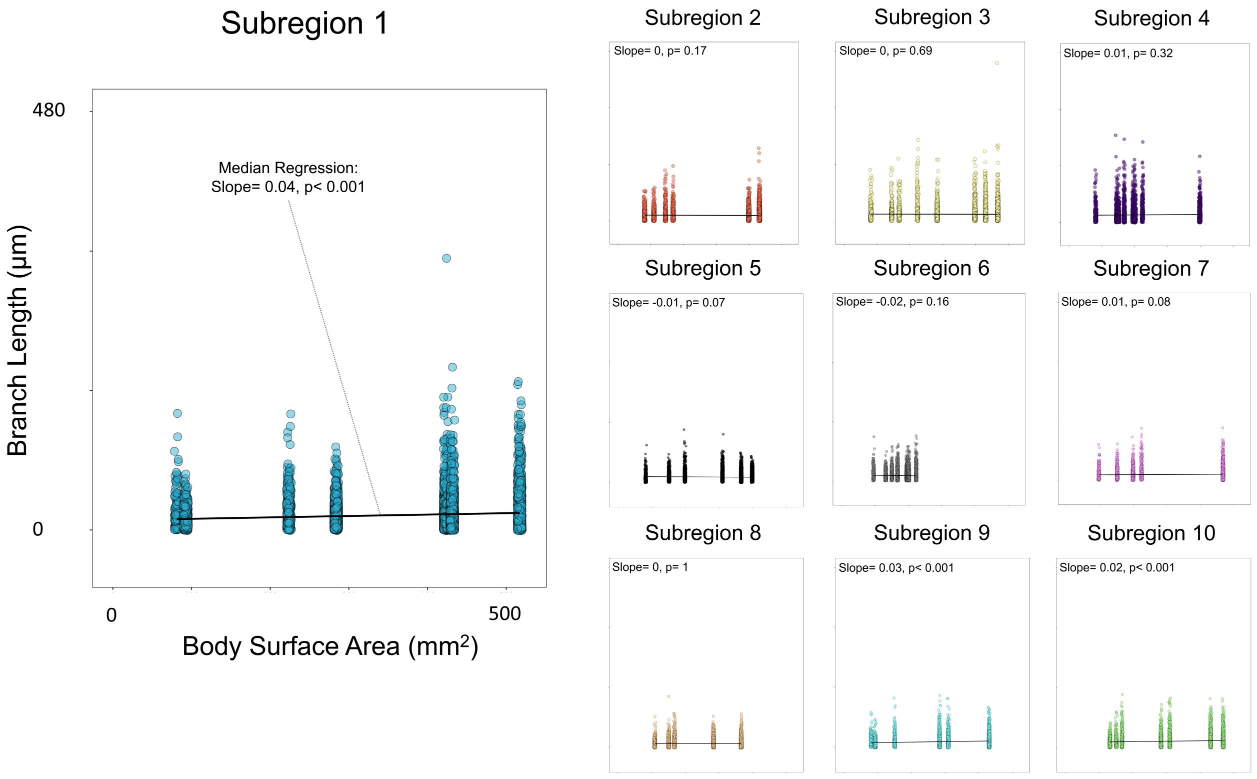


**Supplemental Figure 7. The Relationship Between Branch Length and Body Surface Area.** The relationship between branch length and body surface area was examined. Body surface area was estimated using axial measurements and modelling the shape as an ellipsoid. The relationship was investigated using median regression at each subregion (black line). The slope and p-value are transcribed on each plot. A positive slope denotes longer branches with a larger body size while a negative slope would reveal that the branches are shorter in smaller animals. No relationship was observed between branch length and body size in any of the subregions. This suggests that the number of polygon sides is constant in animals of different sizes. The axes labelling conventions and limits as seen for subregion 1 are the same for all subregions.

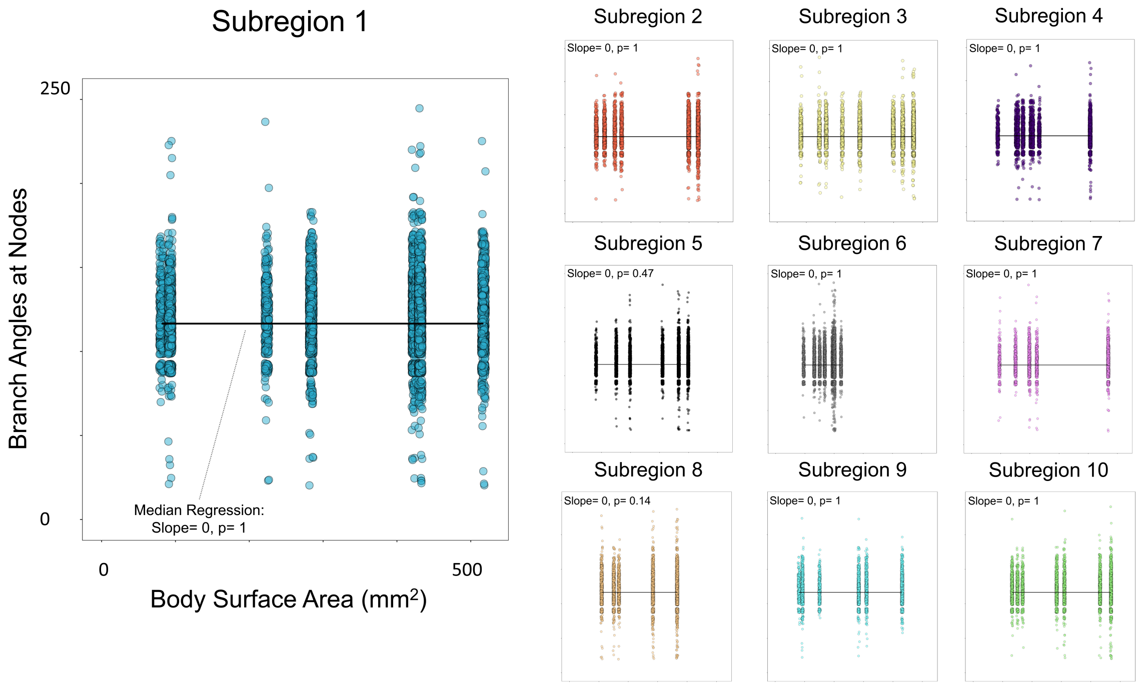


**Supplemental Figure 8. The Relationship Between Branch Angles at Nodes and Body Surface Area.** The relationship between branch angles at nodes and body surface area was examined. Body surface area was estimated using axial measurements and modelling the shape as an ellipsoid. The relationship was investigated using median regression at each subregion (black line). The slope and p-value are transcribed on each plot. A positive slope denotes higher branching angles with a larger body size while a negative slope would reveal that the angles are smaller in smaller animals. No relationship was observed between branch angles and body size in any of the subregions. This suggests that the rules of bifurcation remain constant as an animal grows. The axes labelling conventions and limits as seen for subregion 1 are the same for all subregions.

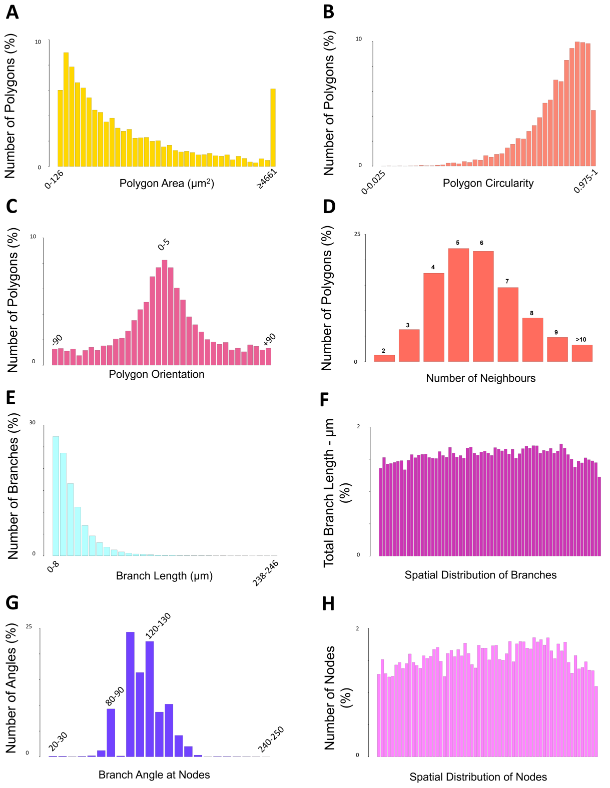


**I**

| Feature | Unit | Number of Bins | Bin Width | First Bin | Last Bin | Bin Upper Limit |
|---|---|---|---|---|---|---|
| Polygon area | μm² | 38 | 126 | 0 | ≥4661 | 15747 |
| Polygon Circularity | NA | 40 | 0.025 | 0 (circle) | ≥0.975 | 1 (line) |
| Polygon Orientation | ° | 36 | 5 | -90 | ≥85 | 90 |
| Number of Polygon Neighbours | NA | 9 | 1 | ≤2 | ≥10 | 16 |
| Branch Length | μm | 31 | 8 | 0 | ≥238 | 246 |
| Spatial Distribution of Branch Length | μm | 64 | The image is divided into a 8x8 grid and each bin represents | NA | Top left of Image | Bottom Right of Image |
| Branch Angle at Nodes | ° | 23 | 10 | 20 | ≥240 | 250 |
| Spatial Distribution of Nodes | NA | 64 | The image is divided into a 8x8 grid and each bin represents | NA | Top left of Image | Bottom Right of Image |

**Supplemental Figure 9. Generating a Feature Vector.** The semi-automated segmentation programme enabled us to extract key morphological characteristics from the nerve net. These features can be visualised in the form of a histogram with defined numbers of bins, bin width and upper/lower limits. The height of a bar relates to the density of a specific parameter. The limit is different for each bin and is detailed in the table (I). (A) Polygon area of entire nerve net population as a histogram. (B) Polygon circularity of entire nerve net population as a histogram. (C) Polygon orientation of entire nerve net population as a histogram. (D) Number of polygon neighbours of entire nerve net population as a histogram. (E) Branch length of entire nerve net population as a histogram. (F) Total branch length within defined windows in analysed regions from the entire nerve net population, displayed as a histogram. (G) Branch angles at nodes of entire nerve net population as a histogram. (H) Number of nodes within defined windows in analysed regions from the entire nerve net population, displayed as a histogram. (I) Table of feature vector histogram parameters. The parameters include number of bins, bin width, final bin threshold and bin upper limit. All features had a constant bin width except, polygon area and number of polygon neighbours in which the last bin included all values above a certain threshold. The spatial distribution features were calculated by dividing each analysed region into an 8x8 grid and measuring the density of the specific feature in that window. The first window is located at the top left of the analysed region, it moves to the right 8 times and then moves down to start again on the left side of the analysed region. This sequence was repeated across the entire area.
